# Identifying Microbial Interaction Networks Based on Irregularly Spaced Longitudinal 16S rRNA sequence data

**DOI:** 10.1101/2021.11.26.470159

**Authors:** Jie Zhou, Jiang Gui, Weston D. Viles, Haobin Chen, Juliette C. Madan, Modupe O. Coker, Anne G. Hoen

## Abstract

The microbial interactions within the human microbiome are complex and temporally dynamic, but few methods are available to model this system within a longitudinal network framework. Based on general longitudinal 16S rRNA sequence data, we propose a stationary Gaussian graphical model (SGGM) for microbial interaction networks (MIN) which can accommodate the possible correlations between the high-dimensional observations. For SGGM, an EM-type algorithm is devised to compute the *L*_1_-penalized maximum likelihood estimate of MIN which employs the classic graphical LASSO algorithm as the building block and can therefore be implemented easily. Simulation studies show that the proposed algorithms can significantly outperform the conventional algorithms when the correlations between measurements grow large. The algorithms are then applied to a real 16S rRNA gene sequence data set for gut microbiome. With the estimated MIN in hand, module-preserving permutation test is proposed to test the independence of the MIN and the corresponding phylogenetic tree. The results demonstrate strong evidence of an association between the MIN and the phylogenetic tree which indicates that the genetically related taxa tend to have more/stronger interactions. The proposed algorithms are implemented in R package *lglasso* at https://CRAN.R-project.org/package=lglasso.

## 1. Introduction

Microorganisms thrive in communities with large numbers. They interact with their host and with one another in various forms such as commensalism, synergism, competition, parasitism, and predation. This complex set of interactions can be depicted in a form of a microbial interaction network (MIN) (Faust and Raes (2012)). Traditionally, such interactions are inferred using culture based methods which can only accommodate small number of microbial strains (Gause (1934); Harcombe (2010)). Since majority of the microbes cannot be cultivated, the estimated interactions in laboratory are of limit and may be misleading. Underpinned by the advances in next generation sequencing (NGS) technologies, a complete microbiome profile can be measured with relatively low cost and therefore provides the opportunity for the researchers to investigate the microbial interaction in real environments. However, the complexities of these high-throughput data such as high dimensionality, zero inflation and compositional nature etc pose substantial challenges for the downstream analysis such as the identification of MIN (Faust and Raes (2012)). Currently, the primary way for the inference of MIN is the so-called pairwise method in which the co-occurrence or mutual exclusion pattern of two species are compared using a measure such as Pearson or Spearman correlation (Arumugam et al (2011); Barberan (2012); Qin et al (2010); Zhou et al (2010)). An emerging method is based on the concept of conditional independence, i.e., the conditional joint distribution of two microbes given all the other microbes in the microbiome. Conditional independence is conceptually superior to the pairwise method since it removes the effects of all the other microbes when measuring the relationship between the given microbes (Viles et al (2021); Chen et al (2017)). Furthermore, if we assume that the data is from a high-dimensional normal distribution, then the inverse matrix of covariance matrix, i.e., so-called precision matrices, directly reflect the conditional independence relationship among microbes. With such an appealing interpretation, precision matrices become the ideal tools to explore the MIN if the assumption of normality is reasonable in real situations.

The estimation of high-dimensional precision matrices themselves has been studied extensively (Wang and Jiang, 2020; Marco et al, 2018), among which the most popular way is the *L*_1_-penalized maximum likelihood estimation. An efficient algorithm called graphical LASSO is proposed to implement the *L*_1_-penalized maximum likelihood estimation of precision matrix (Friedman et al (2008, 2019)). Graphical LASSO estimates the precision matrix recursively based on the well-known coordinate descent algorithm. Another popular yet less efficient method is the so-called neighborhood method (Meinshansen and Buhlmann (2006)). The neighborhood method estimates the neighborhood of each node and then pieces these neighborhoods together to form the final estimate of network. However, one of the prerequisite conditions for these methods is that the data have to be independent and follow the same normal distribution. Though such constraint may be reasonable if the data is cross-sectional, in the clinical trials many data sets are of a longitudinal nature in which multiple observations are made on the same subject. In such studies, the observations such as the gut microbiome from same subject are often correlated and consequently the methods mentioned above can not apply to such situations. It should be noted that there have been studies in literature to estimate the network based on the longitudinal data (Chen et al (2017); Sacha et al (2018)). However, to our knowledge, the majority of the existing studies have employed the time series method, e.g, vector autoregression or state-space model, to deal with the correlation between consecutive observations within the same cluster. Such methods require that the data is equally spaced and the length of the data from a given subject is long enough. These requirements are often violated for many longitudinal data sets in studies of human subjects.

In this paper, we consider the estimation of MIN from irregularly-spaced longitudinal data based on 16S rRNA sequence data. We assume that the relative abundance, after some transformation, are normally distributed and focus on the estimation of its precision matrix. To this end, we first proposed homogeneous stationary Gaussian graphical models (SGGM) in which all the subjects share a same autocorrelation parameter *τ*. For homogeneous SGGM, a recursive graphical LASSO algorithm is proposed to compute the *L*_1_-penalized maximum likelihood estimate (MLE) of the network. Based on this preliminary result, we then propose heterogeneous SGGM in which each subject possesses his/her own autocorrelation parameter *τ*_*i*_. For heterogeneous SGGM, given the complex form of its likelihood function, an expectation-maximization (EM) type algorithm is devised to compute the *L*_1_-penalized MLE of the network and *τ*_*i*_’s. The algorithm integrates the EM algorithm with graphical LASSO and has been implemented in the R package *lglasso*. Simulation studies demonstrate that though the performance of the proposed algorithm is comparable with the existing algorithms such as graphical LASSO (Friedman et al (2008)) or neighborhood method (Meinshansen and Buhlmann (2006)) when the correlations between data are negligible, substantial improvements are achieved by the proposed algorithms when the correlations between data grow large which is exactly what we expected.

In second part, we use the proposed models to study a longitudinal gut microbiome data set from a cohort of children with cystic fibrosis in New Hampshire (Madan et al (2016)). With the estimated MIN in hand, we measured the correlation between the estimated MIN and the corresponding phylogenetic tree. To this end, the length of shortest path between two taxa on the MIN (phylogenetic tree) is used as the measure of their relatedness. Then a module-preserving permutation test procedure is proposed to detect the correlation. The results demonstrate strong evidence for the non-zero correlation between the MIN and phylogenetic tree which indicate that genetically related taxa also have more/stronger interactions. Related discoveries have been noted in other studies (Chaffron et al (2010); Eiler et al (2012)). These findings provide basis for using the phylogenetic tree as a tool to explore the microbial interaction in the future studies (Chung et al (2021)).

The paper is organized as follows. In Section 2, we introduce the main results in which Section 2.1 defines the underlying data generating process stationary Gaussian graphical model (SGGM). Section 2.2 introduces the inference algorithm for the homogeneous SGGM, and Section 2.3 describes the inference algorithm for the heterogeneous SGGM. Section 3 compares the performance of the proposed algorithms with the conventional methods using simulated data. In Section 4, module-preserving permutation test is proposed to explore the relationship between the estimated MIN and phylogenetic tree. We conclude with a brief discussion in Section 5.

## 2. Model and Algorithms

### 2.1. Data generating process

Let 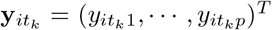 denote observations of some transformed abundance data of microbiome with *p* taxa from subject *i* at time *t*_*k*_ (1 *≤ i ≤ m*, 1 *≤ k ≤ n*_*i*_) so that it is appropriate to assume that 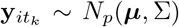 where 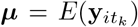 and 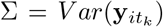 are mean vector and covariance matrix respectively. Precision matrix is defined as Ω = Σ^*−*1^. Then *n*_*i*_*p* vector 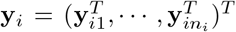 represents all the observations for subject *i*, and vector 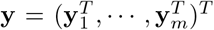 the observations for all the *m* subjects with 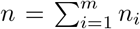. As for the correlation between the observations, we assume that the observations from different subjects are independent, i.e., 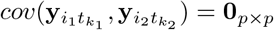 for *i*_1_ ≠*i*_2_, *k*_1_ *≥* 1, *k*_2_ *≥* 1. While for the observations from the same subject, we assume 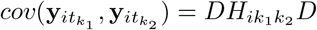 where *D* = diag(*σ*_1_, …, *σ*_*p*_) with 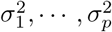the diagonal elements of Σ, while 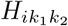 is the correlation matrix between 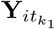 and 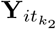 for which the following form is assumed,

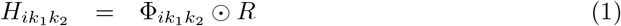

The symbol ⊙ stands for the Hadamard product of matrices 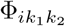and *R*. Here *R* is the correlation matrix of covariance matrix Σ, while matrix 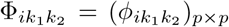 defines the dampening rates at which the absolute values of correlations decrease as time goes from 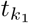 to 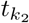. For example, 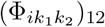 is the dampening rate of the absolute value of correlation 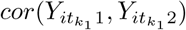 to that of correlation 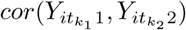. Theoretically, dampening rates can vary from component to component, and also depend on the time points as long as the resulting matrix 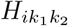 is positive definite. However, in this paper we will assume all the components share the same dampening rate and they depend on time points 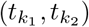only through the distance 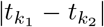, i.e, 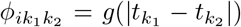 for some decreasing function 0 *≤ g*(*·*) *≤* 1. Motivated by the studies on longitudinal regression model (Diggle et al (2002)), we assume function *g*(*·*) have the form of 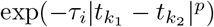. For *p* = 0, we have 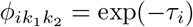 which is called uniform correlation and can be used to model the spatial correlation. For example, specimens may be collected at different body sites from the same subjects for which the uniform correlation seems to be a reasonable assumption. On the other hand, the cases when *p >* 0 can be used to model the irregularly spaced temporal correlation which typically deceases as the time span 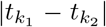increases. In particular, functions 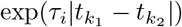 and 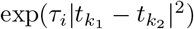have been widely used in longitudinal regression model. Here the parameters *τ*_*i*_’s, which are referred to as autocorrelation parameters, measure the dampening rates that shared by all the components of **y**_*it*_, (*i* = 1, …, *m*). These *τ*_*i*_’s reflect the subject level information which may be used as possible biomarker for some diseases in clinical trial. Without loss of generality, we always employ the correlation function 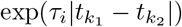 in the following and also assume that the observations have been centered so that ***µ*** = **0**. Let Σ_*i*_ denote the covariance matrix of the observation vector **y**_*i*_, the density function of **y** is then given by

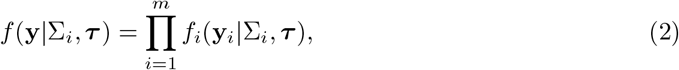

where 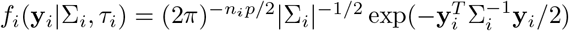 with

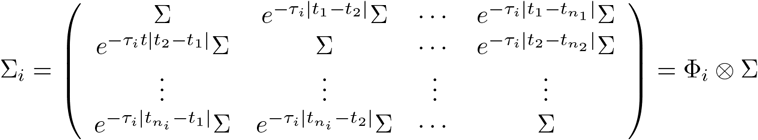

and

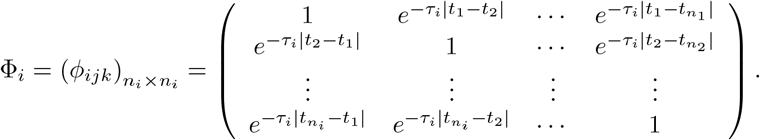

Since 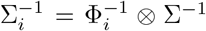 and 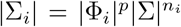, the computation of the inverse matrix of *n*_*i*_*p × n*_*i*_*p* matrix Σ_*i*_ and its determination in density function (2) can be broken down to that of matrix Φ_*i*_ and Σ which can greatly reduce the computation of likelihood function (2). In the following two sections, we consider the selection of Ω and call the model above stationary Gaussian graphical model (SGGM), as opposed to the conventional GGM where the independence between observations 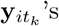 is required. Stationarity stems from the fact that the same Ω is shared by all the subjects and at all time points.

Given the fact that the number of unknown parameters in Ω are much larger than sample size in the context of gut microbiome, the sparsity is typically assumed for the structure of Ω. To this end, *L*_1_-penalized maximum likelihood estimation (MLE) is usually adopted to achieve the sparsity of Ω. If the observations 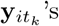 are independent, the graphical LASSO algorithm (Friedman et al (2008)) are the most efficient and popular algorithm to this end. Another popular method is the neighborhood method (Meinshansen and Buhlmann (2006)) which, though easier to be implemented than graphical LASSO, are not the most efficient since it is not based on likelihood function. For correlated data, existing studies focus on the inference of network defined by the state transition matrix between consecutive time points in time series models (Chen et al (2017); Epskamp et al (2018)). Consequently, the times series in these studies have to be long enough and equally spaced. Longitudinal data in human studies often have short length and are irregular spacing and do not satisfy these requirements. With these observations in mind, we propose an algorithm to identify Ω in SGGM (2) which leverages the efficient graphical LASSO to compute the penalized maximum likelihood estimator of precision matrix Ω = Σ^*−*1^. To ease the exposition, an algorithm for so-called homogeneous SGGM are considered first based on which the algorithm for the general SGGM are proposed. Both algorithms yield the penalized likelihood estimates of Ω.

### 2.2. Network with homogeneous SGGM

In this section, we consider identifying Ω under the assumption *τ*_1_ = … = *τ*_*m*_ = *τ* which says that correlations between microbes dampens at the same rate for each subjects in the cohort. From the density function (2), the log-likelihood function for 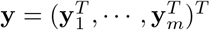 is given by

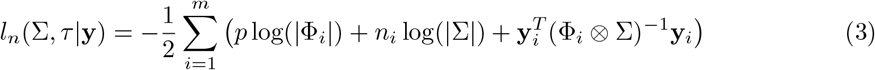

up to a constant. Here we have used the fact that Σ_*i*_ = Φ_*i*_ ⊗ Σ and 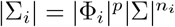. Note that the last term in (3) can be written as,

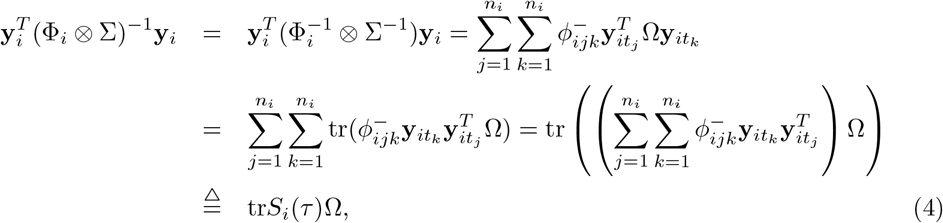

where 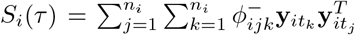 and 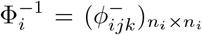. Substituting (4) into (3), we have

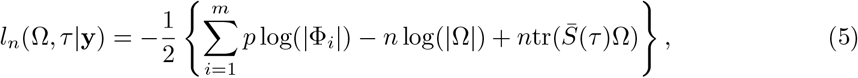

where 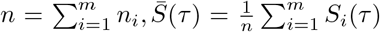. Here we used 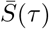to emphasize that matrix 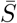 is a function of unknown parameter *τ*. The sparse network can be achieved by minimizing the following *L*_1_-penalized negative log-likelihood function,

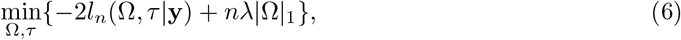

for given tuning parameter *λ >* 0. The minimization problem (6) can be carried out through a block coordinate descent manner. First note for given *τ*, the solution of Ω can be obtained through the following minimization,

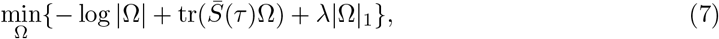

which happens to has the same form as the GGM for independent data when the empirical covariance matrix is given by 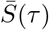.Consequently, graphical LASSO algorithm can be used to compute the sparse estimate of Ω in (7). On the other hand, given Ω, the minimization of (6) with respect to *τ* does not involve any *L*_1_ penalty term and consequently can be carried out through the maximization of likelihood function (5) with respect to *τ*. The conventional Newton algorithm can be used in this step. This process is proceeded until the convergence is achieved. This algorithm will be referred to as homogeneous longitudinal graphical LASSO (LGLASSO) for which the details are summarized in the following table Algorithm 1. Given initial value *τ*_0_ and Ω_0_, tuning parameter *λ* and error tolerance *e >* 0,

#### Algorithm 1 Identify the network for homogeneous SGGM

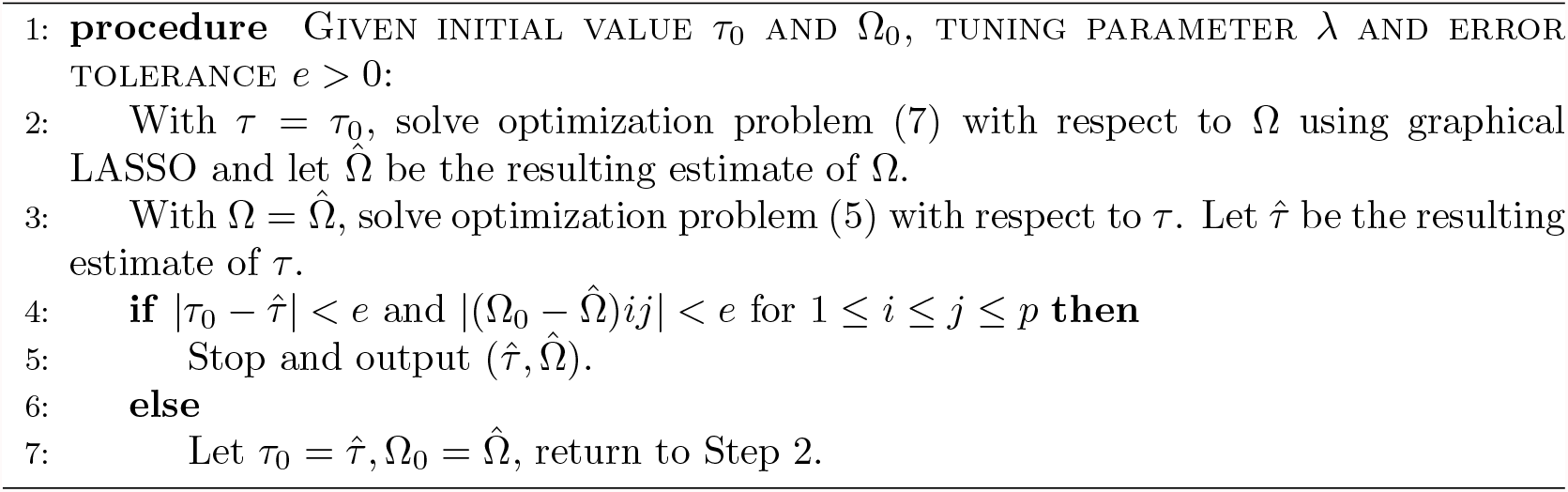

### 2.3. Network with heterogeneous SGGM

In homogeneous SGGM, we have assumed that a single correlation parameter *τ* applies to all the individuals. In real data analysis, this parameter may potentially be affected by some covariates such as gender, diet etc. and also may relate to the outcome. In other words, *τ*’s provide a characterization for each individual. In this section, we consider the network identification without assuming *τ*_1_ = … = *τ*_*m*_. Specifically, we assume the parameters *τ*_*i*_’s are independent random variables with identical exponential distribution, *τ*_*i*_ *∼* exp(*α*). Consequently, the joint density function for 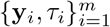 is

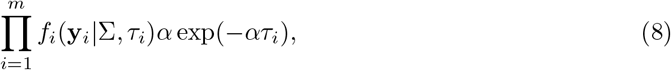

from which the likelihood function is given by

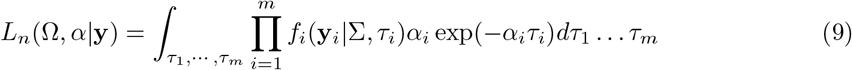

The sparse estimates for the network can then be obtained through minimizing the following *L*_1_-penalized negative log-likelihood function,

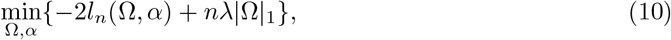

where *l*_*n*_(Ω, *α*) = log(*L*_*n*_(Ω, *α*| **y**)). Since the explicit form of *l*_*n*_(Ω, *α*) is unavailable, Expectation-Maximization (EM) algorithm are proposed here to find the solution to (10) (Dempster et al (1977)). Since we are considering the negative log-likelihood function in (10), the maximization in the EM algorithm will be replaced by minimization here. The correlation parameters ***τ*** = (*τ*_1_, …, *τ*_*m*_) will be taken as the so-called missing data. Recall in the first step of the EM algorithm, the conditional distribution of missing data ***τ*** given **y**, Σ = Σ_0_, *α* = *α*_0_ has to be derived which from (2) and (8) can be shown to be

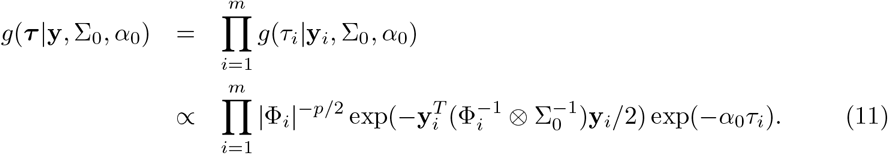

With (11) in hand, the expectation of the complete log-likelihood function for (**y, *τ***) with respect to (11) has to be computed. Given the joint density function 8 of (**y, *τ***), the expectation of its logarithmic transformation can be shown to be

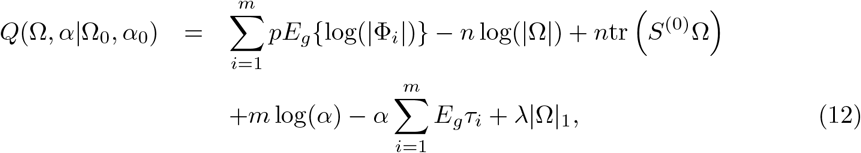

where 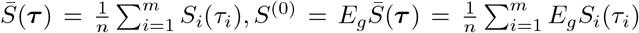 in which *S*_*i*_(·) is defined in Section 2.2. In the second step of the EM algorithm, the minimum of the *Q* function (12) has to be computed. This again is implemented through a block coordinate descent algorithm. First, for fixed Ω, it is straightforward to show that the minimum of *Q* function with respect to *α* is attained at

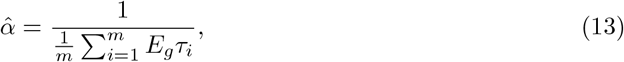

i.e., the reciprocal of the sample mean of the conditional expectation of *τ*_*i*_ with respect to density (11). Then, for given *α*, the minimization of (12) with respective to Ω is equivalent to

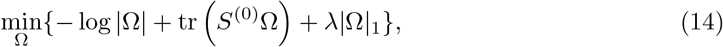

which can also be solved through the graphical LASSO algorithm. The difficult part of this algorithm is to find the expectation 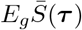, which may not admit an explicit form given the fact that 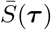 is a nonlinear function of ***τ*** and the complex form of density function *g*(***τ***); therefore this expectation will be computed through a Monte Carlo method. This algorithm will be referred to as heterogeneous longitudinal graphical LASSO for which the details are summarized in the following table Algorithm 2. Given initial value *α*_0_ and Ω_0_, tuning parameter *λ* and error tolerance *e >* 0,

### Remark

(1) Both algorithms 1 and 2 leverage the graphical LASSO algorithm to achieve their efficiency even though graphical LASSO itself is devised for the independent data. This stems from the observation that graphical LASSO does not rely on the data directly; instead, it relies on a positive-definite matrix *S* involved in the objective function. For independent data, *S* is just the covariance matrix; while in the context of longitudinal data, it is *S*^(0)^ defined above; (2) For a given sequence of tuning parameters *λ*’s, a solution path can be generated by the proposed algorithms. In order to select the optimal network Ω from this model pool, a variety of model selection criteria can be used, e.g, Akaike information criterion (AIC), Bayesian information criterion (BIC) or cross validation (CV) etc. In the following numerical studies, we use the extended BIC (EBIC) to select the tuning parameter (Foygel and Drton (2010); Zhou et al (2021)).

#### Algorithm 2 Identify the network for heterogeneous SGGM

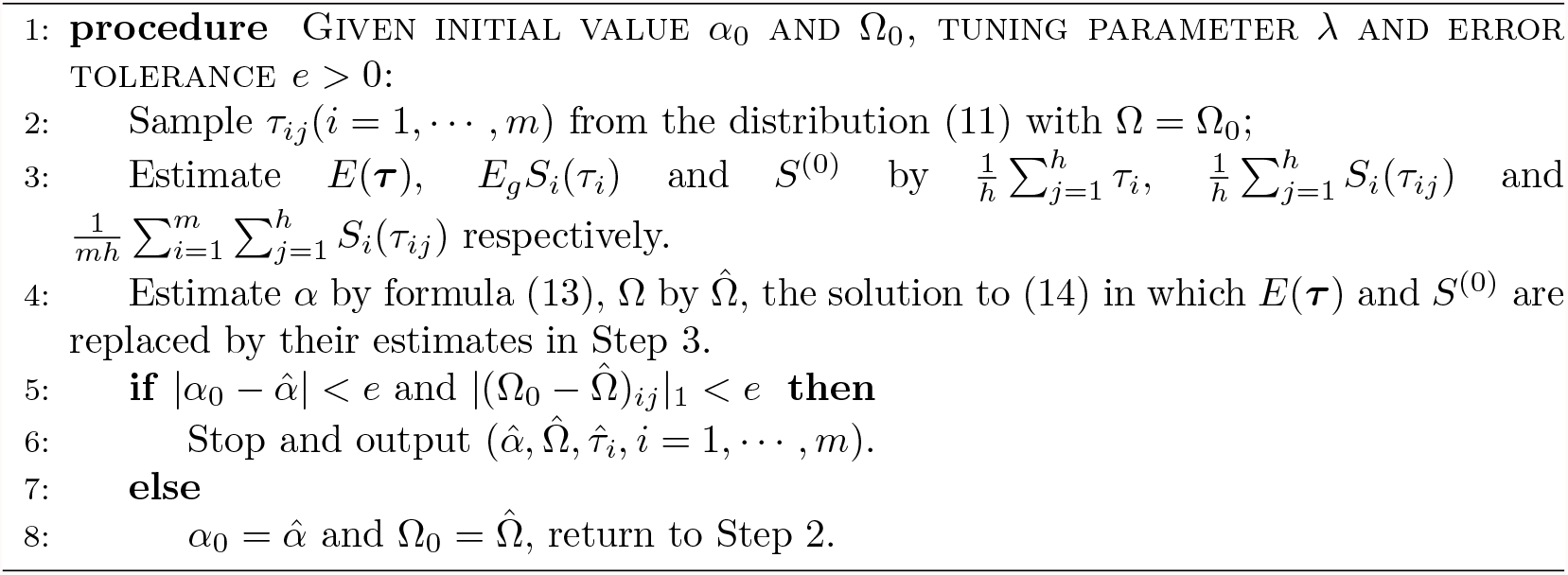

## 3. Simulation

In this section, we compare the proposed algorithm with conventional network selection methods, such as graphical LASSO (Friedman et al (2008, 2019)) and neighborhood algorithm (Meinshansen and Buhlmann (2006)) under different parameter settings. We demonstrate that the proposed longitudinal graphical LASSO algorithms can outperform the existing algorithms when the correlations between the observations within the same cluster increase. Motivated by indices such as true positive rate (TPR) and false positive rate (FPR), the following two indices are defined to measure the difference of two given network *G*_1_ and *G*_2_ which share the same nodes,

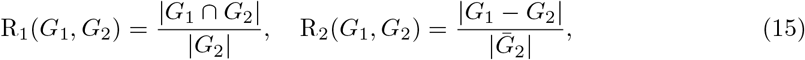

where |*G*_2_| represents the number of edges in 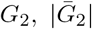 the number of null edges in *G*_2_, |*G*_1_ *∩G*_2_| the number of shared edges by *G*_1_ and *G*_2_ and |*G*_1_ *−G*_2_| the number of edges that appear in *G*_1_ while not in *G*_2_. It is straightforward to show that in case of *R*_1_ = 1 and *R*_2_ = 0, these two network *G*_1_ and *G*_2_ just share the same edge set, i.e., *G*_1_ = *G*_2_. Particularly, when *G*_2_ is the underlying true network and *G*_1_ represents the estimated network, then R_1_ and R_2_ are just the conventional TPR and FPR respectively.

First we show that the networks estimated by longitudinal graphical LASSO coincide with the networks estimated by graphical LASSO when the autocorrelation parameter between the observations within the same cluster decreases to zero. Specifically, we consider a network with *p* = 100 nodes that is shared by *m* = 20 subjects. The precision matrix corresponding to this network is generated through the R package *BDgraph* with edge density equal to 0.2. For each subject, observations at four time points are generated. For subject *i*, the space between two consecutive time points *t*_*ij*_ and *t*_*i*(*j*+1)_ is generated by max*{*Δ_*ij*_, 0.5*}* where Δ_*ij*_’s follow a Poisson distribution with mean value 1. Then the data for subject *i* is generated with real autocorrelation parameter *τ*_0_ = 0.5. For this data set, both longitudinal graphical LASSO and graphical LASSO are employed to estimate the underlying network. Let *G*_1_ and *G*_2_ denote the respective resulting network from these two algorithms for a given tuning parameter *λ*.

Figure 1 plots the indices *R*_1_ and *R*_2_ as functions of autocorrelation parameter *τ* based on 50 replicates with tuning parameter *λ* = 0.1. Recall that larger *R*_1_ and smaller *R*_2_ means that the two networks bear more similarities. It can be seen from Figure 1 that for large *τ*, the difference between *G*_1_ and *G*_2_ is small. In particular, for *τ >* 3, there is little difference between *G*_1_ and *G*_2_. However, for small *τ*, these two algorithm can generate rather different network. This phenomenon is expected since from equation (2), as *τ* becomes large, the SGGM that is used to model the data converges to the GGM and so they should have smaller difference with larger *τ* .

**Figure 1:**
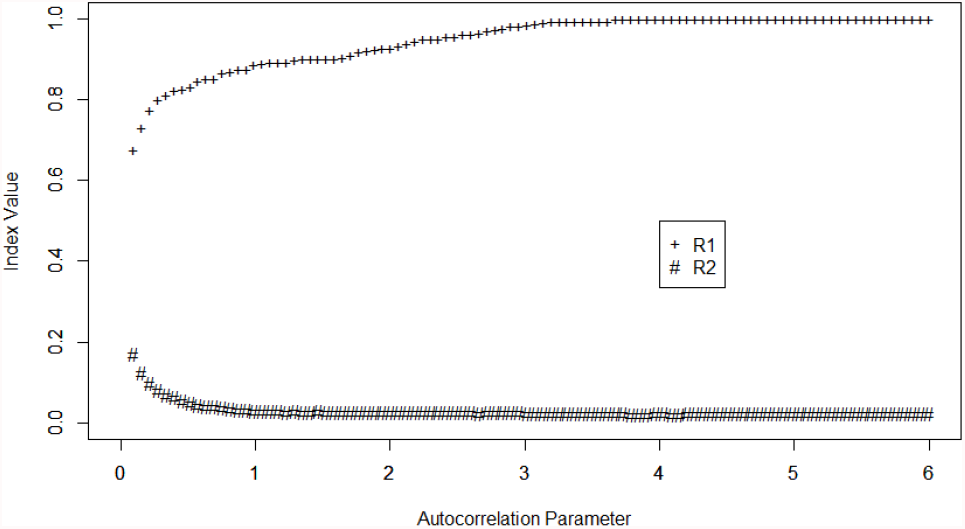
Comparison of longitudinal graphical LASSO (homogeneous) and graphical LASSO. Two indices *R*_1_ and *R*_2_ are used to measure their differences which are the functions of *τ*. Tuning parameter *λ* is set to be 0.1.

In the following, we compare the performance of longitudinal graphical LASSO, graphical LASSO and neighborhood method based on networks with *p* = 80 nodes. Both homogeneous and heterogeneous versions of the algorithm are considered with EBIC as the model selection criterion.

For the homogeneous model, the numbers of subjects considered are *m* = 15, 30 respectively. In each case, the numbers of time points are set to be 7 and 14 respectively. Three choices of correlation parameter are *τ* = 0.1, 0.4, 0.8 respectively. For each combination of these parameters, the data is generated by the same procedure as in Figure 1. The indices TPR and FPR are used to measure the performances of the algorithms. The results for the three algorithms are listed in Table 1. From Table 1, it can be seen that for majority of the cases considered, the proposed longitudinal graphical LASSO has the lowest FPRs and highest TPRs. Another observation is that neighborhood method seems to outperform graphical LASSO and be more robust with regard to the correlation between the data. This may be attributed to the fact that, contrary to graphical LASSO, neighborhood method does not rely on the likelihood function of the data directly and therefore less impacted by the correlation between the data.

**Table 1:** Performance comparison of homogeneous longitudinal graphical LASSO (lglasso), graphical LASSO (glasso) and neighborhood method (nh) for network selection.

| | | $\tau = 0.1$ | | $\tau = 0.4$ | | $\tau = 0.8$ | |
| --- | --- | --- | --- | --- | --- | --- | --- |
| glasso |  | TPR | FPR | TPR | FPR | TPR | FPR |
| $m = 15$ | $n_i = 7$ | 0.622 | 0.207 | 0.672 | 0.150 | 0.715 | 0.123 |
| $m = 15$ | $n_i = 14$ | 0.744 | 0.258 | 0.785 | 0.164 | 0.783 | 0.113 |
| $m = 30$ | $n_i = 7$ | 0.758 | 0.257 | 0.795 | 0.160 | 0.807 | 0.119 |
| $m = 30$ | $n_i = 14$ | 0.776 | 0.206 | 0.798 | 0.087 | 0.772 | 0.045 |
| nh |  | TPR | FPR | TPR | FPR | TPR | FPR |
| $m = 15$ | $n_i = 7$ | 0.628 | 0.132 | 0.702 | 0.122 | 0.745 | 0.115 |
| $m = 15$ | $n_i = 14$ | 0.678 | 0.125 | 0.729 | 0.082 | 0.754 | 0.062 |
| $m = 30$ | $n_i = 7$ | 0.738 | 0.122 | 0.776 | 0.084 | 0.731 | 0.056 |
| $m = 30$ | $n_i = 14$ | 0.716 | 0.099 | 0.721 | 0.041 | 0.747 | 0.022 |
| lglasso |  | TPR | FPR | TPR | FPR | TPR | FPR |
| $m = 15$ | $n_i = 7$ | 0.752 | 0.075 | 0.737 | 0.075 | 0.723 | 0.068 |
| $m = 15$ | $n_i = 14$ | 0.802 | 0.066 | 0.796 | 0.065 | 0.792 | 0.061 |
| $m = 30$ | $n_i = 7$ | 0.824 | 0.071 | 0.814 | 0.069 | 0.801 | 0.067 |
| $m = 30$ | $n_i = 14$ | 0.875 | 0.040 | 0.869 | 0.041 | 0.865 | 0.041 |

For the heterogeneous model, the number of subjects considered is set to be 20. The numbers of time points are set to be *n*_*i*_ = 10, 20, 30 respectively. The parameter *α* in exponential distribution are set to be *α* = 1.25, 2.5, 10 respectively, which approximately correspond to the *τ*’s with mean values 0.8, 0.4, 0.1 respectively in Table 1. For each of these *α*’s, 20 random samples are generated form the corresponding exponential distribution which will be used as the real autocorrelation parameters for the 20 subjects. The spaces between consecutive time points are generated using the same procedure as in Figure 1. From the heterogeneous longitudinal graphical LASSO, we obtain two types of results. One is the estimated network and the others are the predictions of the 20 *τ*_*i*_’s. Table 2 lists the results for the estimated network from longitudinal graphical LASSO, graphical LASSO and neighborhood method respectively. It can be seen that heterogeneous longitudinal graphical LASSO outperforms the other two for the majority of cases considered, with highest TPRs and lowest FPRs. As for the prediction of *τ*_*i*_, we consider two indices to evaluate the performance of predictors. One is the average relative error (ARE) which is defined as 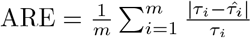. The other is the correlation between the true *τ*’s and the predicted value 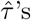. The results for these two indices are listed in Table 3. From Table 3, we can see that, for a given *α*, ARE decreases while correlation increases as sample size increases. In particular, the estimates of correlation parameter is strongly correlated with the true values of *τ*_*i*_. Such correlations turn out to be rather robust to the choice of tuning parameter of *λ*. In other words, as a subject level index, the predicted 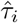 reflects the real characteristic of subject *i* very well.

**Table 2:** Performance comparison of heterogeneous longitudinal graphical LASSO (lglasso), graphical LASSO (glasso) and neighborhood method (nh) for network selection.

| | | $\alpha = 1.25$ | | $\alpha = 2.50$ | | $\alpha = 10$ | |
| --- | --- | --- | --- | --- | --- | --- | --- |
| glasso |  | TPR | FPR | TPR | FPR | TPR | FPR |
| $m = 20$ | $n_i = 10$ | 0.667 | 0.133 | 0.678 | 0.194 | 0.632 | 0.242 |
| $m = 20$ | $n_i = 20$ | 0.695 | 0.114 | 0.726 | 0.149 | 0.633 | 0.196 |
| $m = 20$ | $n_i = 30$ | 0.744 | 0.116 | 0.714 | 0.177 | 0.708 | 0.206 |
| nh |  | TPR | FPR | TPR | FPR | TPR | FPR |
| $m = 20$ | $n_i = 10$ | 0.737 | 0.128 | 0.725 | 0.167 | 0.671 | 0.175 |
| $m = 20$ | $n_i = 20$ | 0.790 | 0.101 | 0.788 | 0.135 | 0.745 | 0.208 |
| $m = 20$ | $n_i = 30$ | 0.821 | 0.084 | 0.753 | 0.101 | 0.733 | 0.182 |
| lglasso |  | TPR | FPR | TPR | FPR | TPR | FPR |
| $m = 20$ | $n_i = 10$ | 0.779 | 0.128 | 0.813 | 0.167 | 0.837 | 0.169 |
| $m = 20$ | $n_i = 20$ | 0.756 | 0.052 | 0.810 | 0.072 | 0.851 | 0.118 |
| $m = 20$ | $n_i = 30$ | 0.801 | 0.038 | 0.815 | 0.053 | 0.873 | 0.073 |

**Table 3:** Evaluation of the predicts of *τ*_*i*_’s in heterogeneous longitudinal graphical LASSO. Here ARE stands for the average relative error, and Cor stands for the correlation between the true *τ*_*i*_’s and their estimates 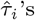.

| | | $\alpha = 1.25$ | | $\alpha = 2.5$ | | $\alpha = 10$ | |
| --- | --- | --- | --- | --- | --- | --- | --- |
|  |  | ARE | Cor | ARE | Cor | ARE | Cor |
| $m = 20, n = 10$ | | 0.235 | 0.980 | 0.253 | 0.982 | 0.453 | 0.965 |
| $m = 20, n = 20$ | | 0.133 | 0.987 | 0.135 | 0.992 | 0.232 | 0.987 |
| $m = 20, n = 30$ | | 0.105 | 0.992 | 0.109 | 0.995 | 0.164 | 0.993 |

## 4. Gut Microbial Interaction Network and Phylogenetic Tree

In this section, a longitudinal data set from a cohort of children with cystic fibrosis was investigated using the proposed heterogeneous longitudinal graphical LASSO in Section 2. Specifically, the stool samples from thirty-eight children were collected during the period of 6 month to 51 months old (Madan et al (2016)). The number of observations from each child ranges from 2 to 17. Each observation consisted of the abundances of 16,383 amplicon sequence variants (ASV) for the 16S rRNA gene. These sequences were then collapsed to the genus level taxa using R package *DADA2* (Callahan et al (2016)). The sequences that have no genus level information were dropped. Then all the taxa with the proportion of nonzero observations less than 10% were combined together which was referred to as composite taxon. This left us 83 taxa in total. The observations of zeros for each of these 83 taxa were replaced by the minimum abundance of that taxon divided by 10. The log-ratio transformation was then carried out for the relative abundance in which the composite taxon was taken as the reference. Application of the heterogeneous model in Section 2.3 to the transformed data generated the final estimated network which is displayed in Figure 2. The corresponding predicted correlation parameters *τ*_*i*_’s are displayed in Figure 3. Based on the modularity maximization algorithm (Newman (2006); Vincent et al (2008)), eight communities were identified in the estimated network which are listed in Table 4. This step was accomplished through R package *igraph* (Csardi (2006)).

**Table 4:**
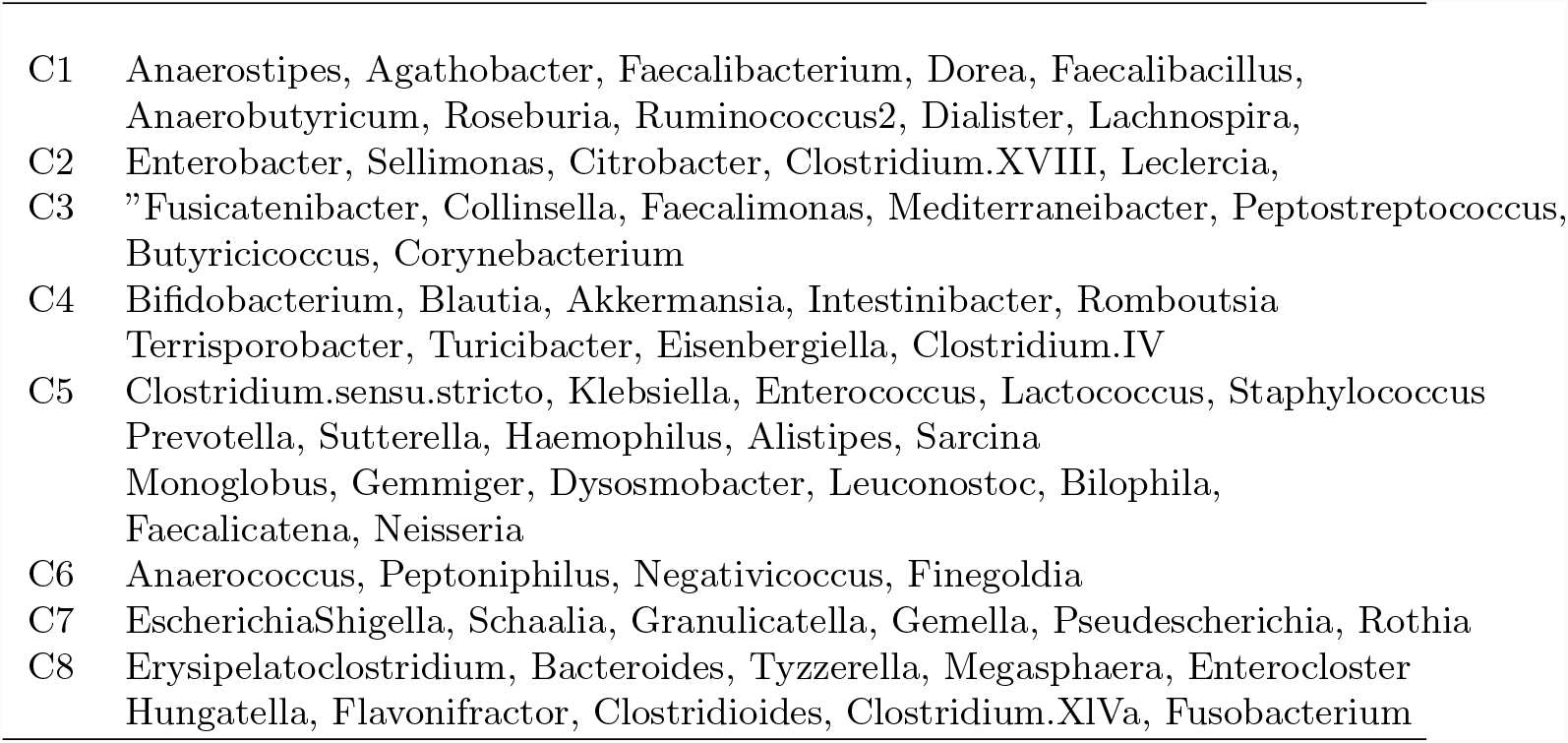
Eight communities selected by maximizing the modularity of the estimated MIN in Figure 2.

**Figure 2:**
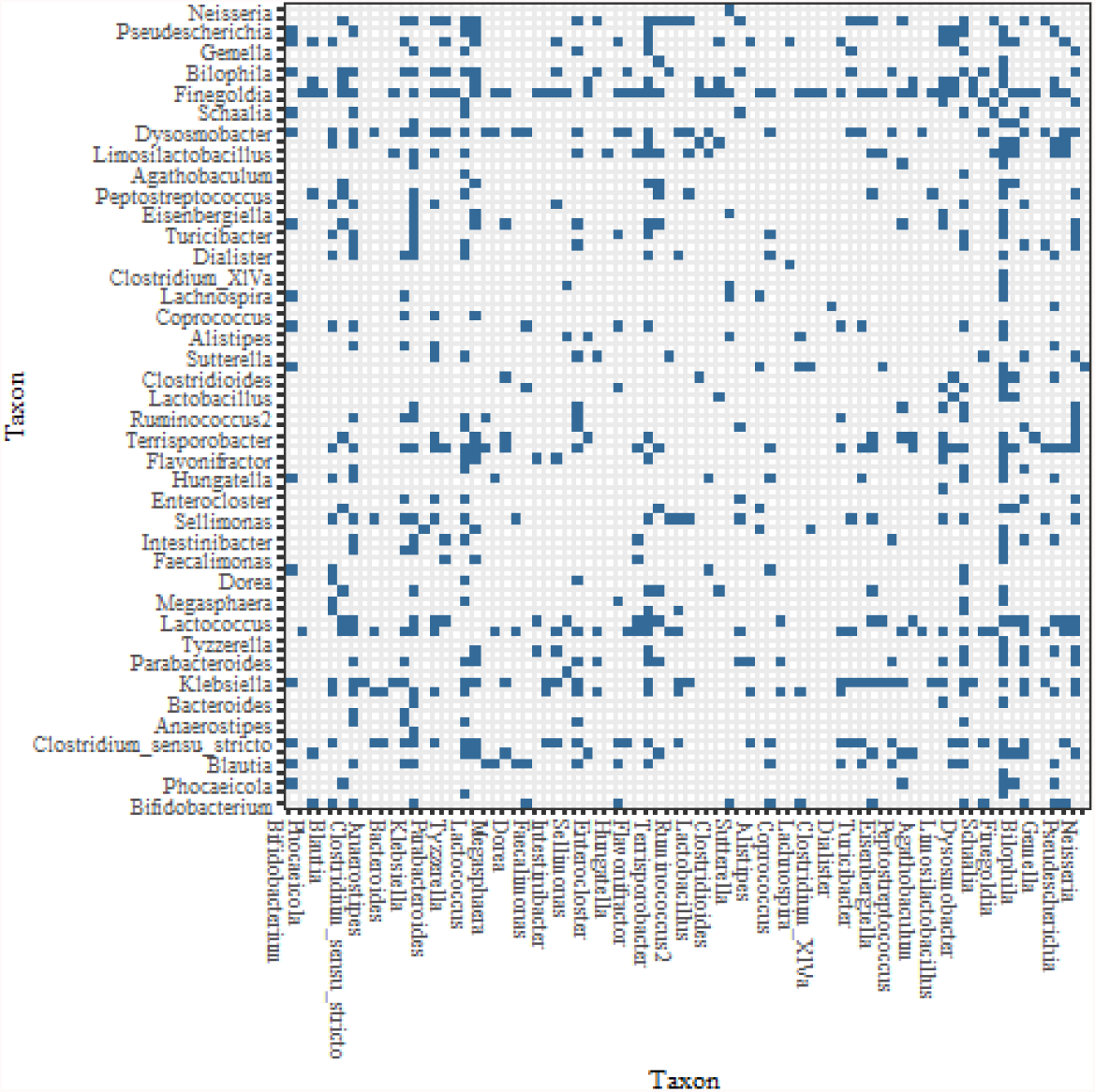
Microbial interaction network generated by heterogeneous longitudinal graphical LASSO based on the longitudinal data set in Section 4. A dark dot represents an edge between the two microbes labeled on its x and y coordinate. Only half the microbes are labeled for the sake of clarity. The complete information are included in the supplementary material in form of adjacent matrix.

**Figure 3:**
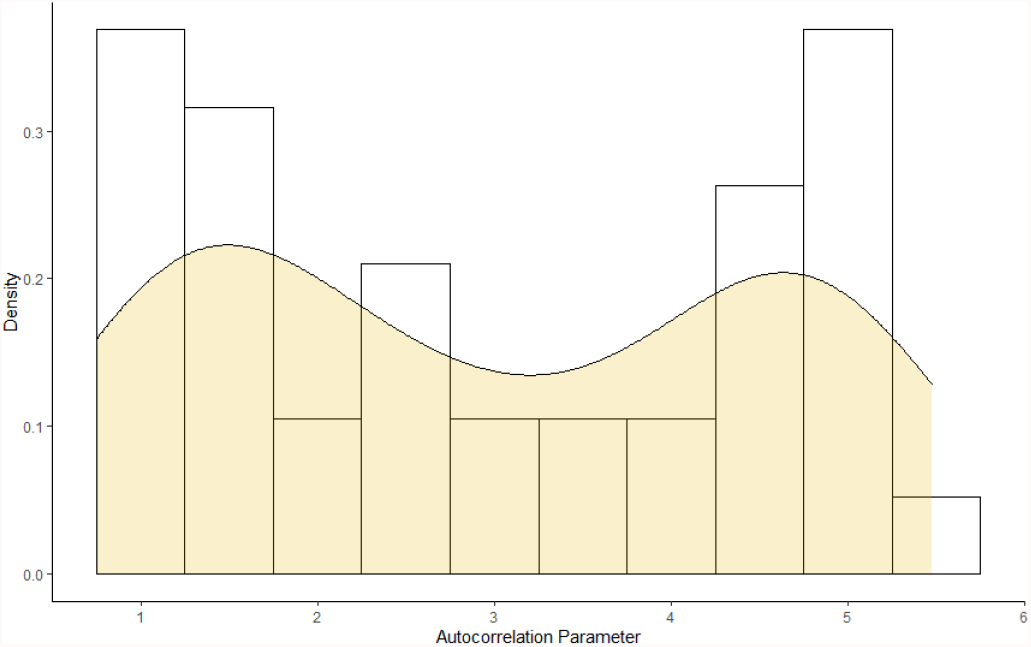
Histogram for the individual estimated autocorrelation parameter 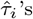 for 38 subjects in the longitudinal data set in Section 4.

In order to demonstrate the estimated network can reveal the true structure of the underlying network, we investigate the relationship between this estimated network and phylogenetic tree of the 82 taxa (the composite one is excluded here). The pholygenetic tree constructed from the same data set is presented in Figure 4. Our underlying theory is that microbial taxa that are closer in evolution histories also have more/stronger interactions in human body. To validate this theory, we set the null hypothesis as the estimated network being independent with the phylogenetic tree. To test this hypothesis, we compute the distance between two taxa in both the estimated network and phylogenetic tree. Here the distance between taxa A and B is defined as the length of the shortest path from A to B in the estimated network or phylogenetic tree. Let *d*_1_, *d*_2_ be the distance variables from the estimated network and phylogenetic tree respectively, then the correlation between *d*_1_ and *d*_2_ is used to measure the relatedness between the network and phylogenetic tree. This correlation turns out to be *r*_0_ = 0.244 for the tuning parameter selected by EBIC. Five thousand bootstrap samples yields the 95% confidence interval of *r*_0_ to be [0.213, 0.275], see Figure 5.

**Figure 4:**
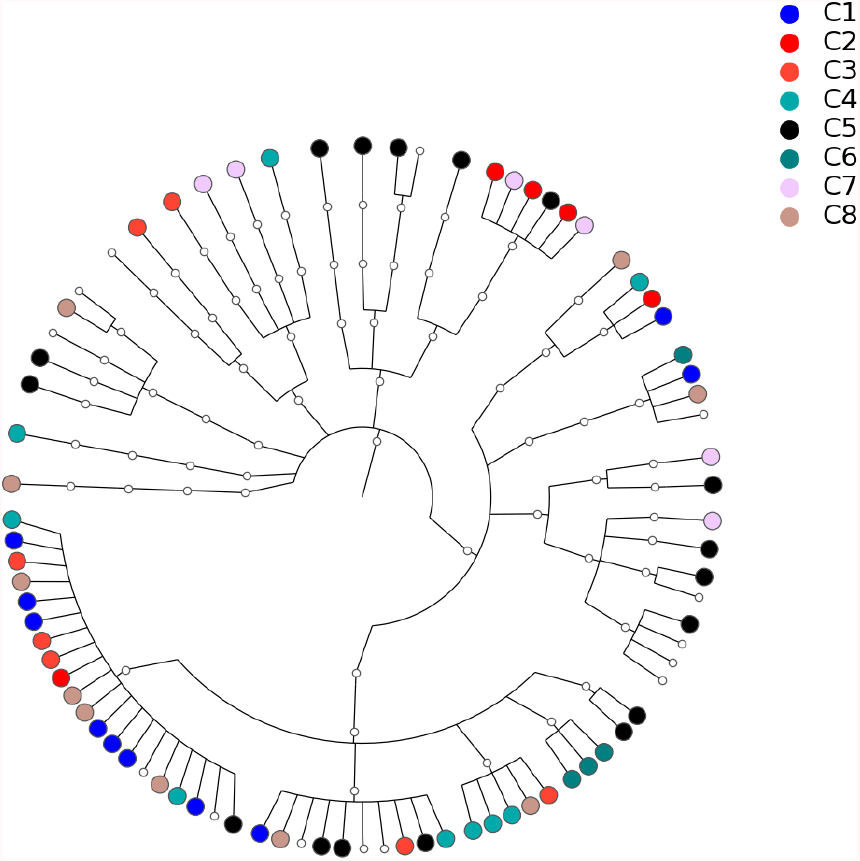
Phylogenetic tree for the 82 microbial taxa in the longitudinal data set in Section 4. Dots with different color correspond to different communities listed in Table 4.

**Figure 5:**
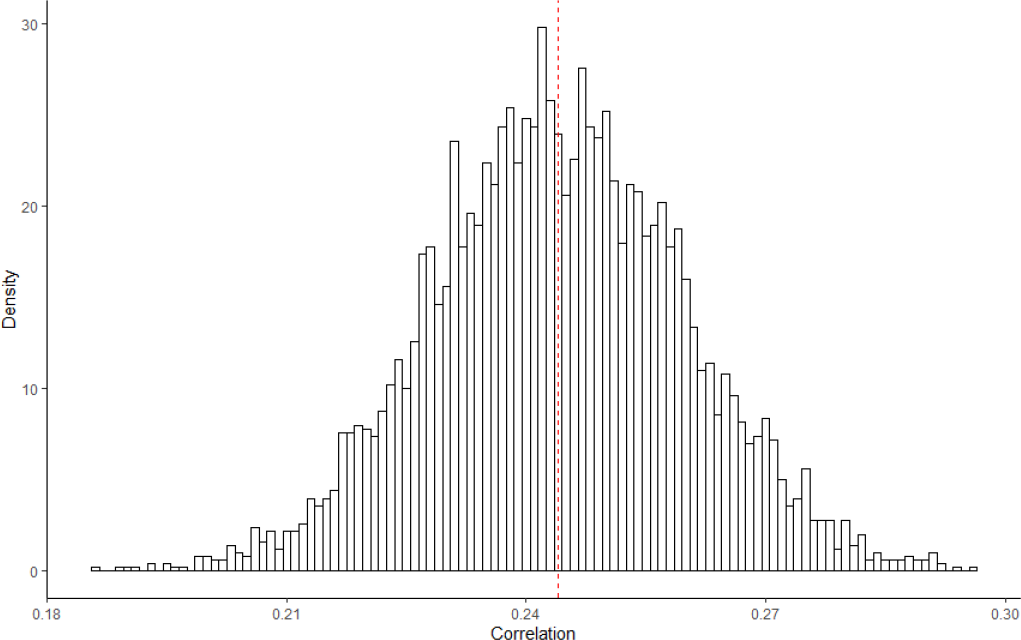
Five thousand bootstrapped correlations between the estimated MIN in Figure 2 and the phylogenetic tree in Figure 4. The red dashed line represents the mean value.

In order to understand the significance of *r*_0_, we use permuted data to empirically estimate the null distribution. Out of multiple ways to carry out the permutation test, we propose a module-preserving permutation test here. Specifically, we keep the structure of the estimated network unchanged, and permute the order of the 82 taxa *m* = 5000 times. For the *i*th permuted network, distance vector 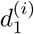 is computed from which correlations 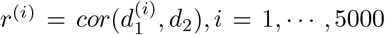 can be derived. Note that the studies in Diciccio and Romano (2017) show that it is not appropriate to use these permuted correlation *r*^(*i*)^ directly to determine the significance of *r*_0_ since it can lead to misleading p-value and the so-called large type III error rate. Here we adopt the proposal in Diciccio and Romano (2017) and use the following more robust studentized correlation, 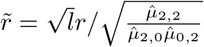, where 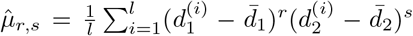, *l* is the length of vector *d*_1_(*d*_2_) and 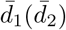 are the mean value of *d*_1_(*d*_2_). Based on 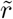, the p-value of 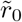 turns out to be smaller than 1/5000, which means that the correlation between the estimated network and phylogenetic tree is statistically significant, i.e., the data supports the theory that microbial taxa with closer evolution histories also tend to have more/stronger interactions.

It should be noted that some related findings have been reported in other studies. For example, Chaffron et al (2010) found that genomes from coexisting microbes tended to be more similar than expected by chance, both with respect to pathway content and genome size. The studies in Eiler et al (2012) also revealed that ecological coherence was often dependent on taxonomic relatedness. However, it should be pointed out that those studies have employed the correlation coefficient as the basis to infer the relatedness of microbes which may lead to misleading conclusion. It is well known that the correlation between two microbes A and B may be induced by the correlation of these two microbes with another microbe C, even though A and B will be independent if C is fixed. In current study, the relatedness between the microbes are defined based on the conditional correlation coefficient which by its definition eliminates the possible spurious correlation between microbe A and B induced by microbe C. Therefore, our conclusion that MIN and phylogenetic tree are significantly related is more convincing and stronger than the existing findings in literature.

Note the results above are derived with the tuning parameter selected by EBIC. The correlations between MIN and phylogenetic tree are actually pretty robust with respect to the tuning parameter. To demonstrate this point, let us consider the networks generated from other choices of tuning parameter. Specifically, for each of 40 tuning parameters ranging from *λ* = 45 to *λ* = 55, the same heterogeneous model and the permutation test are carried out. The estimated and permuted correlations are displayed in Figure 6 in the form of boxplot. The boxplots in Figure 6, from left to right, correspond to the tuning parameters increasing from 45 to 55 respectively. The dots linked by the line represent the correlation between the estimated MIN’s and phylogenetic tree while others correspond to the correlations computed from the permuted MINs. The most prominent feature of Figure 6 is that in almost all of the 40 cases, the observed studentized correlations between MIN and phylogenetic tree are positive and significant. The boxplots in the middle part of the plot demonstrate more variability which also correspond to larger real correlation coefficients.

**Figure 6:**
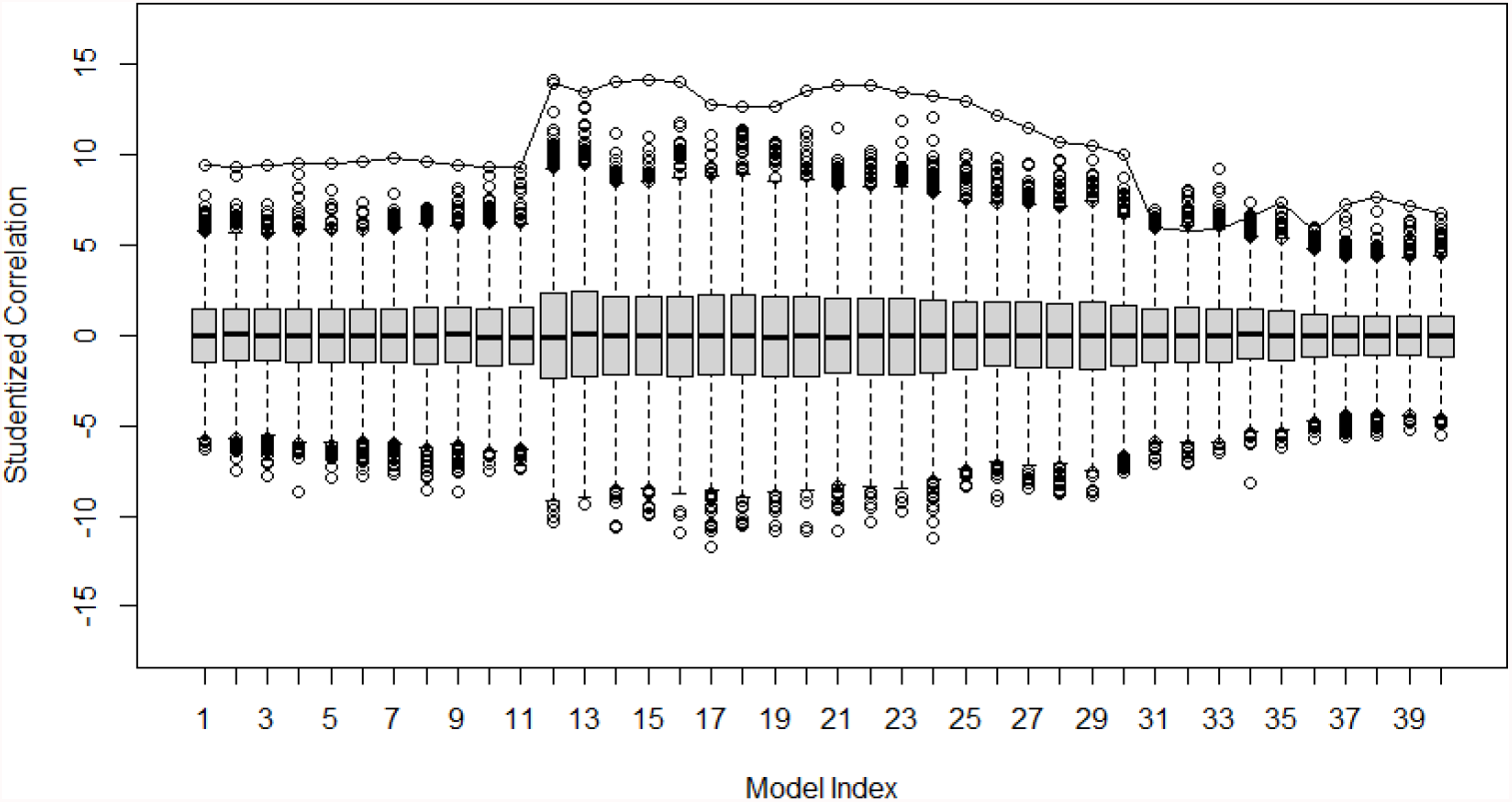
Forty boxplots of correlations between the estimated networks and phylogenetic tree which correspond to, from the left to right, the tuning parameters *λ* increasing from 45 to 55. For a given boxplot, the dots linked are the estimated correlations from the original estimated network; others are correlations computed from 5000 permutations of MIN.

In addition, we also considered the tuning parameters smaller than 45 or larger than 55, the maximum correlations are still achieved in the same region as Figure 6. In summary, the correlation between the selected MIN and phylogenetic tree is not only statistically significant, but also higher than the correlations derived based on most of other tuning parameters. This partly validates that EBIC is a reasonable model selection method for microbial interaction network.

## 5. Conclusion

We proposed algorithms to estimate microbial interaction networks based on irregularly spaced longitudinal 16S rRNA gene sequence data. These algorithms show their advantages over the traditional methods where the correlations are omitted or handled using traditional time series model. For the estimated network, a module-preserving permutation test was designed to test its effectiveness, which discovered the relationship between microbial interaction network and phylogenetic tree and validated the proposed algorithms. It should be noted that we have assumed that the microbes of interest in given subject share the same time dampening rate. Though it seems to work well for the 16S rRNA gene sequence data of gut microbiome, there may be occasions in which different taxa need different time dampening rates. Identifying ways to expand the proposed algorithms to such situations will be the subject of future work.

## Acknowledgments

This work is supported in part by National Natural Science Foundation of China (90308210044), US National Institutes of Health grants (R01LM012723).

## Notes

https://github.com/jiezhou-2/lglasso

https://CRAN.R-project.org/package=lglasso

### Competing Interest Statement

The authors have declared no competing interest.

### Summary of Updates

(1)Correct the typos and grammatical errors in the first version; (2)Simulation, Real data analysis are more accessible;

